# Identification of pathogens in culture-negative infective endocarditis cases by metagenomic analysis

**DOI:** 10.1101/336388

**Authors:** Jun Cheng, Huan Hu, Yue Kang, Weizhi Chen, Wei Fang, Kaijuan Wang, Qian Zhang, Aisi Fu, Shuilian Zhou, Chen Cheng, Qingqing Cao, FeiYan Wang, Shela Lee, Zhou Zhou

## Abstract

Pathogens identification is critical for the proper diagnosis and precise treatment of infective endocarditis. Although blood and valve cultures are the gold standard for IE pathogens detection, many cases are culture-negative, especially in patients who had received long-term antibiotic treatment, and precise diagnosis has therefore become a major challenge in the clinic. Metagenomic sequencing can provide both information on the pathogenic strain and the antibiotic susceptibility profile of patient samples without culturing, offering a powerful method to deal with culture-negative cases. In this work, we assessed the feasibility of a metagenomic approach to detect the causative pathogens in resected valves from IE patients.

Using our in-house developed bioinformatics pipeline, we analyzed the sequencing results generated from both next-generation sequencing and Oxford Nanopore Technologies MinION nanopore sequencing for the direct identification of pathogens from the resected valves of seven clinically culture-negative IE patients according to the modified Duke criteria. Moreover, we were able to simultaneously characterize respective antimicrobial resistance features. This provides clinicians with valuable information to diagnose and treat IE patients after valve replacement surgery.

## Introduction

Infective endocarditis (IE) is a serious disease associated with significant morbidity and mortality(1-3), whose prognosis strongly depends on early diagnosis and optimized antibiotic therapy. Therefore, identifying the underlying pathogens responsible for IE is critical. Currently, blood and valve cultures are the gold standard for IE pathogens detection, but they are time-consuming and infeasible for fastidious or intracellular microorganisms(4), which is a major clinical problem. Although targeted amplicon sequencing such as 16S rRNA sequencing overcomes the limitations of conventional culture-based methods, it can only be used to screen for bacteria(5,6) and does not provide any antibiotic susceptibility information.

Rapid advancements in sequencing technologies provide us with new tools for microbial identification without the need for culturing(7-9). The feasibility of direct pathogens identification from IE samples by short-read whole-genome sequencing on next-generation sequencing (NGS) platforms has been demonstrated in several studies(10,11). Recently, an increased number of studies have shown promise for metagenomics analysis using nanopore long-read sequencing in the rapid detection of microorganisms in clinical samples, including virus from blood samples and bacteria from urine samples(12-14)

To evaluate the analytical and clinical sensitivity and specificity of metagenomics analysis in IE diagnosis, we analyzed the sequencing results generated from both NGS and nanopore sequencing in this study. Sequencing platform-specific bioinformatics pipelines were designed and developed in-house to identify pathogens and detect antimicrobial resistance (AMR) in seven culture-negative IE patients.

## Materials and Methods

### Sample collection and information

The resected valves were collected from the Center of Cardiac Surgery in Fuwai Hospital, National Center for Cardiovascular Diseases (Beijing, China), from April 2017 to August 2017. The study was approved by the ethics committee of the hospital. All patients involved in this study provided their written informed consent, and samples were used for research only. In our study, we included seven patients (six men and one woman, Table S1). These patients were all diagnosed with definite IE (D.IE) according to the modified Duke criteria. The specimens were cut into two equal-sized pieces using sterile scissors in a biosafety cabinet. One piece of tissue was randomly selected for immediate culturing, while the other was snap-frozen at −80°C for metagenomic sequencing and Sanger validation.

### Valve culture (VC) and blood culture (BC)

The specimens were physically ground into particles using a sterile grinder, then placed in sterile tubes containing 5 ml of brain-heart infusion broth and incubated in a CO_2_ enriched atmosphere (5%) at 35°C for 7 days. Growth was evaluated daily. After 7 days of incubation, all samples were subcultured onto blood agar plates (Oxoid, Beijing, China), chocolate agar plates (Oxoid) and MacConkey agar plates (Oxoid), regardless of whether or not growth was suspected. An average of three sets of blood samples were drawn by peripheral venous puncture prior to antibiotic use. Blood samples (about 10 ml for adults, 1–3 ml for children) were injected into aerobic and anaerobic blood culture bottles (Becton Dickinson, Sparks, MD, USA). Blood culture bottles were then loaded into an automated continuous monitoring system (BD BACTEC™ FX400, USA) within 1 h of being drawn and were incubated at 35°C for 7 days. If the subculture of the blood or valves showed bacterial growth, identification was carried out by VITEK MALDI-TOF mass spectrometry (bioMérieux, Marcy l’étoile, France) and antibiotic susceptibility testing was performed subsequently with VITEK 2 COMPACT (bioMérieux).

### DNA extraction and NGS with BGISEQ-500

The frozen valves were thawed at room temperature for 30 min and were then cut into pieces as small as possible with sterile scissors. Approximately 25 mg of tissue was treated with proteinase K (No.148012595, Qiagen, Hilden, Germany) before DNA extraction. Total DNA was extracted using a TIANamp Micro DNA kit (DP316, Tiangen Biotech, Beijing, China) according to the manufacturer’s recommendation. The extracted DNA was fragmented with a Bioruptor (ThermoFisher Scientific, Waltham, MA, USA) instrument to generate 200–300 bp fragments. Libraries were then prepared as follows: first, the DNA fragments were subjected to end-repair and A-tailing; second, the resulting DNA was ligated with bubble-adapters that contained a barcode sequence, and then amplified with PCR. Quality control was carried out with an Agilent 2100 (Agilent Technologies, Santa Clara, CA, USA) to assess the fragment size and using a Qubit dsDNA HS Assay kit (ThermoFisher Scientific) to measure the DNA library concentrations. Qualified libraries were pooled together to form single-stranded DNA (ssDNA) circles and then DNA nanoballs were generated with rolling circle replication. The final DNA nanoballs were loaded onto a sequencing chip and were sequenced with a BGISEQ-500 platform (BGI-Tianjin). Human sequence data were excluded by mapping to a human reference (hg19) using the Burrows–Wheeler alignment tool. After removing human sequences, the remaining sequencing data were aligned to four microbial genome databases, consisting of viruses, bacteria, fungi and parasites. The mapped data were processed for advanced data analysis. We downloaded the latest version of the microbial reference genomes from NCBI (ftp://ftp.ncbi.nlm.nih.gov/genomes/). Currently, our databases cover 1,428 bacterial species, 1,130 viral species related to human diseases, 73 fungal species related to human infections, and 48 parasites associated with human diseases. We used the SOAP Coverage software from the SOAP website (http://soap.genomics.org.cn/) to calculate the multi-parameters of the species.

### PCR and Sanger validation

Extracted DNA of IE resected valves was simultaneously validated by Sanger sequencing, using specific PCR primers: 5%-AGAGTTTGATCCTGGCTCAG-3% and 5%-GGTTACCTTGTTACGACTT-3%. PCR reactions were performed as follows: 96°C for 150 s; (96°C, 30 s; 55°C, 30 s, and 72°C, 90 s) for 30 cycles, then 72°C for 7 min, ending at 4°C. PCR products were detected by agarose gel electrophoresis and purified with a gel extraction kit (DC3511-02, Biomiga Inc., San Diego, CA, USA). Sanger sequencing was performed on an ABI PRISM 3730 DNA Sequencer (Applied Biosystems, Foster City, CA, USA) for validation. Finally, the sequences were analyzed for IE pathogens identification by alignment with sequences in the NT database using the NCBI Blast online software (http://blast.ncbi.nlm.nih.gov/Blast.cgi?PROGRAM5blastn&PAGE_TYPE5BlastSearch&LINK_LOC5blasthome).

### MinION library preparation and sequencing

The frozen valves were thawed at room temperature for 30 min and were then cut into pieces as small as possible with sterile scissors. Approximately 25 mg of tissue was treated with proteinase K before DNA extraction. Total DNA was extracted using QIAamp DNA Mini Kit (Cat No. 51304, Qiagen) according to the manufacturer’s recommendation. Library preparation was performed using the Ligation Sequencing Kit (SQK-LSK108) and Native Barcoding Kit (EXP-NBD103) for genomic DNA, according to the standard 1D Native barcoding protocol provided by the manufacturer (Oxford Nanopore). Briefly, 1.2 μg of extracted genomic DNA from each resected valve sample was fragmented with g-TUBE (Covaris) at 5,000 rpm for 1 min. To perform end-repair, 45 μL of fragmented DNA was mixed in a 0.2 ml PCR tube with 3 μL of Ultra II End-prep enzyme mix (New England BioLabs, NEB), 7 μL of Ultra II End-prep reaction buffer (NEB), and 5 μL of nuclease-free water. The mixture was incubated at 20°C for 5 min, then at 65°C for 5 min. Next, 500 ng of end-prepped samples were combined with 2.5 μL of Native Barcode (one barcode per sample) and 25 μL of Blunt/TA Ligase Master Mix. The mixtures were incubated at 21°C for 30 min.

A total of 700 ng of barcoded libraries were pooled together with 20 μL of Barcode Adapter Mix (BAM) and 10 μL of Quick T4 DNA ligase was added. The mixture was incubated for 10 min at room temperature. The constructed library was loaded into the Flow Cell R9.4 or R9.5 (FLO-MIN106 or FLO-MIN107) of a MinION device, which was run with the SQK-LSK108_plus_Basecaller script of the MinKNOW1.7.14 software.

### Quality control analysis of the NGS data and nanopore data

From the pair-end 150 bp sequence data generated from the BGI platform, low-quality reads, adapter contamination, and duplicated reads and short reads (length <35 bp) were removed. The remaining sequences were then used in further analysis. For the sequencing data obtained from the Nanopore MinION sequencer, base-calling tools in Albacore were used to base-call the data in fast5 files and de-multiplex the data to fastq files for each sample. After quality control analysis, reads with lengths longer than 500 bp and mean quality scores >6 were used in further analysis.

### Species identification of pathogens in seven clinical samples using NGS data and nanopore data

For species identification, first reads originating from the host genome were depleted. In detail, after quality control analysis, reads were aligned with the human genome GRCh38.p11 using bwa mam in the BWA software (genome download from ftp://ftp.ncbi.nlm.nih.gov/genomes/all/GCA/000/001/405/GCA_000001405.26_GRCh38.p11). Reads that could not be mapped to the human genome were retained and aligned with the microorganism genome database for pathogens identification. Our microorganism genome database contained genomic sequences from 259 bacteria, 5,591 fungi and 236 viruses, and sequences from 47 plasmids (plasmid sequences are from ftp://ftp.ncbi.nlm.nih.gov/genomes/refseq/plasmid, and other sequences are from ftp://ftp.ncbi.nlm.nih.gov/genomes/all/). A k-mer alignment algorithm named Centrifuge(15) was used to identify the pathogens in each sample. Species with identified reads ⩽2 for nanopore data and ⩽10 for NGS data were removed, and for those remaining, the relative enrichment rate by query length was calculated and normalized according to genome size. Species with a relative enrichment rate >20% were reported, whereas species with a relative enrichment rate >0.2% and <20% were analyzed further by sampling 200 reads to verify the identify accuracy by blastn(16)in the NT database. Verified species were reported. Finally, all species in the report list were re-calculated for their relative enrichment rate.

### AMR detection among the identified IE pathogens using NGS and nanopore data

After species identification, reads that could not be mapped in the human genome were used for AMR analysis. Species identification tags were added and reads were aligned in the AMR database CARD(17)by Blastn. For all query results, hits with blast e-values <e^−30^ were picked for further analysis. For AMR gene tracking, when sequences were aligned, if hits were lacking in the 5′ or 3′ regions of the gene but coverage of the central part of the gene was observed that would be sufficient to be reported as an AMR gene. For the nanopore data, because of the long read lengths, support from one read was acceptable, but support from three reads was needed for the NGS data. For AMR SNP sites, the coverage level for the gene in which the SNP was located was required to be the same as that from which the AMR gene was detected. Furthermore, each SNP site required support from more than two reads for the nanopore data and three reads for the NGS data. After data had been obtained for AMR genes and SNP sites, the results were organized by drug resistance type using the annotation in the CARD database. Finally, species identification tags were used to map AMR genes to the species level.

## Results

### Clinical characteristics and diagnosis of seven IE patients

To assess the feasibility of metagenomic analysis in the identification of IE pathogens, seven IE patients were included in this study, with most of these patients being male (n=6, 85.7%) with a mean age of 48.3 (Table S2). Our strategy was to employ NGS and nanopore sequencing-based metagenomics analysis to identify IE pathogens with verification provided by Sanger sequencing and traditional clinical diagnosis methods (Fig 1 and Fig 2).

**FIG 1:**
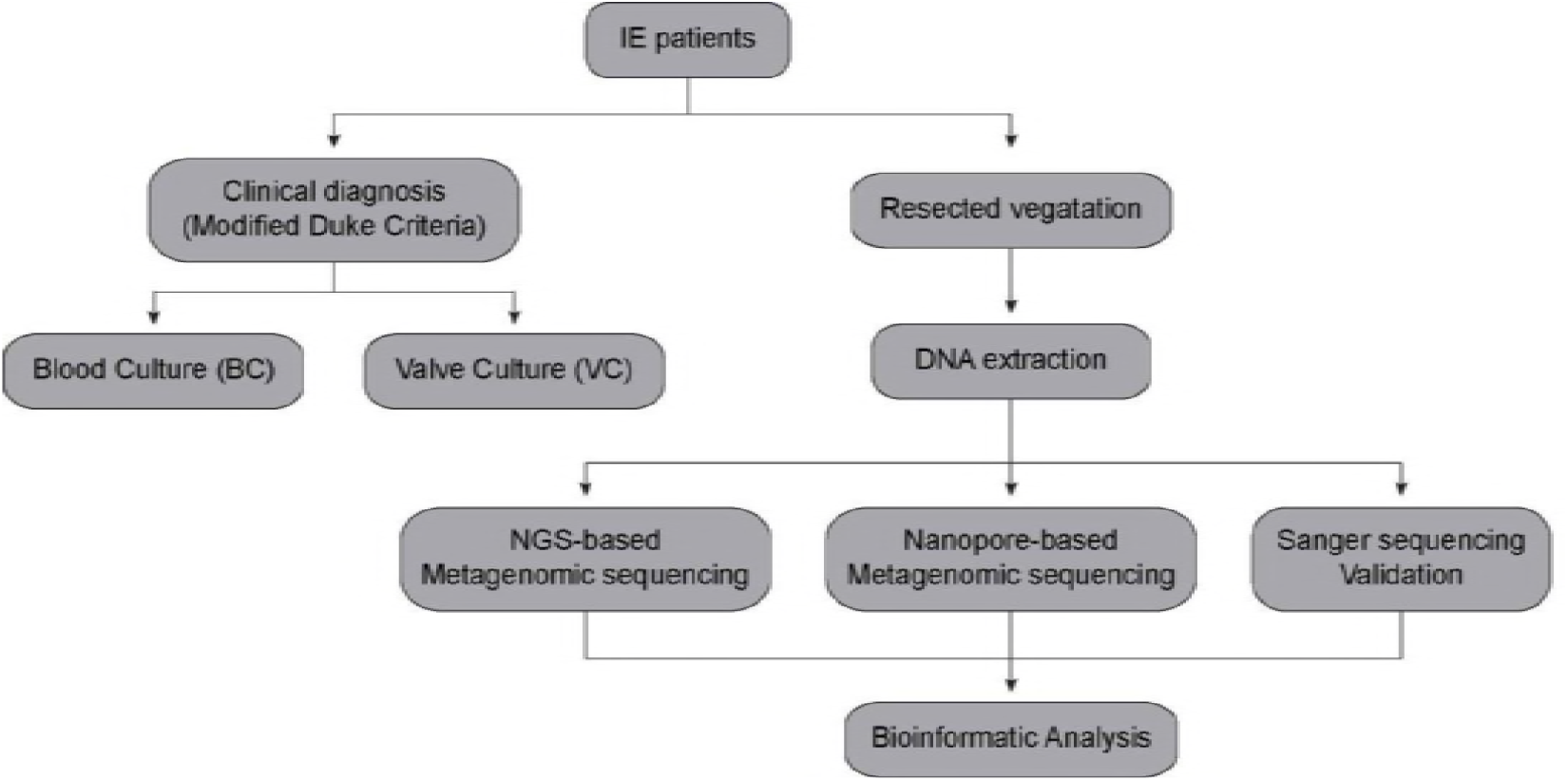
Workflow of IE patient diagnosis with traditional clinic methods and sequencing methods.

**FIG 2:**
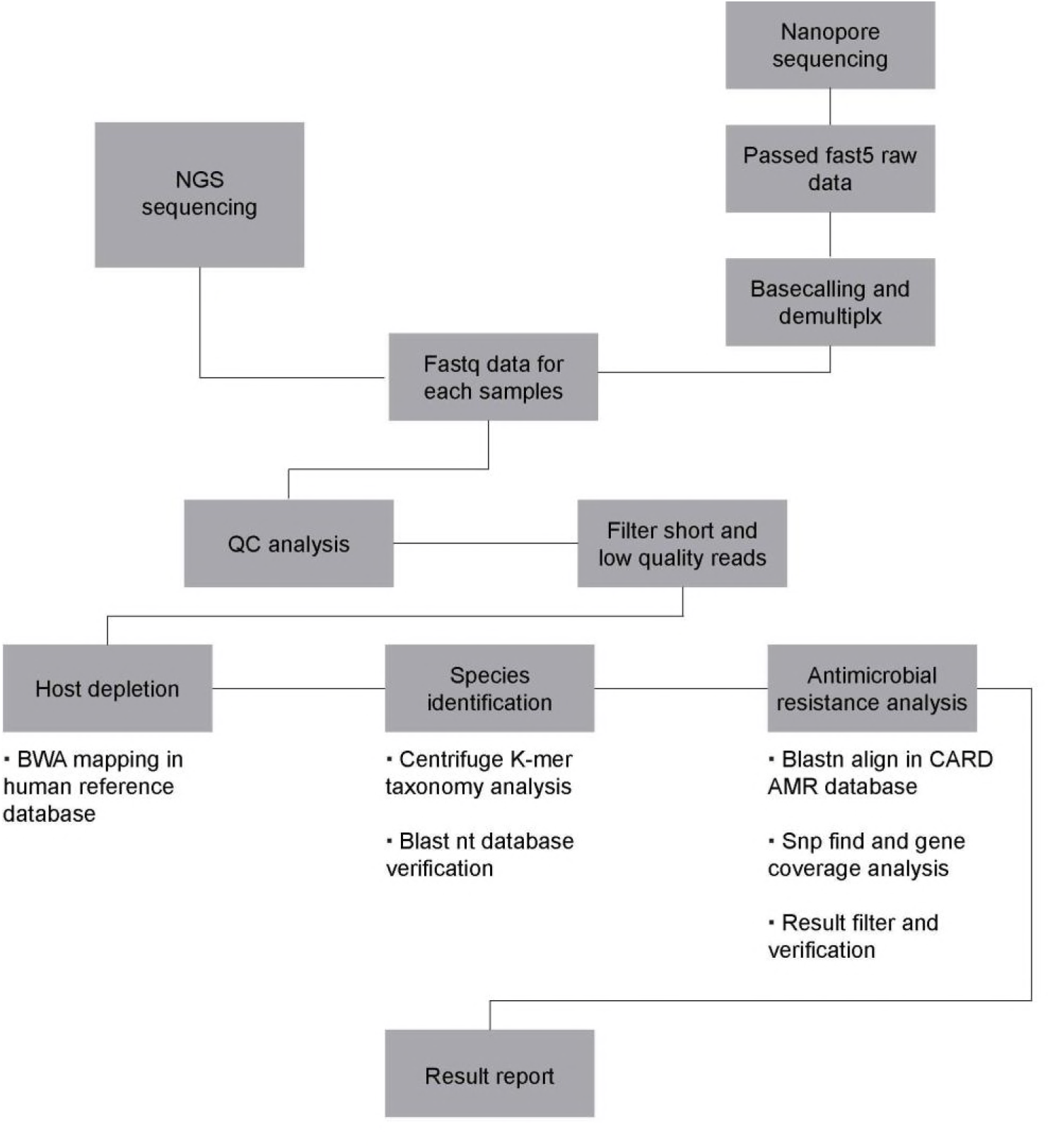
The bioinformatics pipeline for NGS and nanopore sequencing metagenomic analysis.

The patients were firstly scheduled for systemic examinations in the hospital and all were clinically diagnosed as definite cases of IE according to the modified Duke criteria (Fig 1 and Table 1). Most of the blood culture results were negative (n=5) except for *Streptococcus oralis* detected in patient A5 and *Streptococcus anginosus* detected in patient A7 (Table 1). Valve replacement surgeries were then performed and the resected valves were used for Gram-staining and culturing. All of the valve culture results were negative except for one, which was considered to be due to contamination (Table 1).

**TABLE 1.** Clinical diagnosis and Main laboratory results.

|  | Diagnosis | Valve<br>gram<br>staining | Blood<br>culture | Valve<br>culture | Nanopore | NGS | Sanger |
| --- | --- | --- | --- | --- | --- | --- | --- |
| A1 | Definite<br>IE<sup>a</sup> | GPC<sup>b</sup> | negative | *Filamentous*<br>*fungi*<sup>c</sup> | *S. gordonii* | *S. gordonii* | *S. gordonii* |
| A2 | Definite<br>IE<sup>a</sup> | GPC<sup>b</sup> | negative | negative | *S.oralis* | *S.oralis* | *S.viridans*<br>*spp* |
| A3 | Definite<br>IE<sup>a</sup> | negative | negative | negative | *Coxiellaburnetii* | *Coxiellaburneti* | *Not*<br>*detected* |
| A4 | Definite<br>IE<sup>a</sup> | negative | negative | negative | *Bartonella*<br>*Quintana* | *Bartonella*<br>*Quintana* | *Bartonella*<br>*Quintana* |
| A5 | Definite<br>IE | negative | *S.oralis* | negative | *S.oralis* | *S.oralis* | *S.viridans*<br>*spp* |
| A6 | Definite<br>IE<sup>a</sup> | GPC<sup>b</sup> | negative | negative | *S.sanguis* | *S.sanguis* | *S.sanguis* |
| A7 | Definite<br>IE | GPC<sup>b</sup> | *S.anginosus* | negative | *S.anginosus* | *S.anginosus* | *S.anginosus* |

<sup>a</sup>Definite IE was diagnosed according to histopathologic examination, clinical presentation and echocardiographic result.
<sup>b</sup>GPC, gram positive coccus.
<sup>c</sup>This result was considered to be contamination.

### NGS-based metagenomic analysis for the detection of IE pathogens

Resected valves were then used for metagenomics analysis based on NGS. The total DNA of each patient’s valve was extracted and then fragmented to generate 200– 300-bp fragments, which were used to construct a library according to the manufacturer’s protocol (BGI-Tianjin, Tianjin, China; see details in the Materials and Methods section). The final library was sequenced using the BGISEQ-500 platform to generate sequencing data.

After analyzing the data for quality control, the remaining fastq reads for each sample were collected with data volumes of 4.1G (A1), 17G (A2), 3.3G (A3), 4.4G (A4), 8.8G (A5), 3.1G (A6), and 6G (A7). These data were then subjected to bioinformatic analysis to detect pathogen species and AMR genes (see details in the Materials and Methods section).

Metagenomic analysis of the NGS data generated reads of the possible IE pathogens detected for all seven samples (4,260 reads of *Streptococcus gordonii* for A1, 25,275 reads of *S. oralis* for A2, 3,921 reads of *Coxiella burnetii* for A3, 29,438 reads of *Bartonella quintana* for A4, 54,881 reads of *S. oralis* for A5, 370 reads of *Streptococcus sanguinis* for A6, and 45,880 reads of *S. anginosus* for A7) (Table 2). Other information such as pathogen coverage and the depth of the NGS sequencing data were also analyzed (Fig 3A, S1A, and Table 2). Because the AMR profile of an IE pathogens provides valuable information that can guide treatment, a specific bioinformatics pipeline was developed to detect the AMR genes present in these bacteria (Fig 2 and Table 3, S3).

**TABLE 2.** Detail of the results for pathogen species identification from NGS (BGI) data.

| Sample ID | Pathogen species | Genome size | Reads num | Unique reads num | Relative abundance | Coverage | Depth |
| --- | --- | --- | --- | --- | --- | --- | --- |
| A1 | Streptococcus gordonii | 2196662 | 4465 | 4260 | 82.40% | 21.33% | 1.4380 |
| A2 | Streptococcus oralis | 1958690 | 31754 | 25275 | 81.01% | 68.50% | 3.5609 |
| A3 | Coxiella burnetii | 1995488 | 4014 | 3921 | 100.00% | 20.32% | 1.1890 |
| A4 | Bartonella quintana | 1581384 | 29676 | 29438 | 99.55% | 77.68% | 2.9408 |
| A5 | Streptococcus oralis | 1958690 | 68435 | 54881 | 81.74% | 75.74% | 6.8056 |
| A6 | Streptococcus sanguinis | 2388435 | 380 | 370 | 86.20% | 2.33% | 1.0434 |
| A7 | Streptococcus anginosus | 2233640 | 47829 | 45880 | 87.82% | 61.85% | 5.1198 |

**TABLE 3.** AMR analysis results from two different platform sequencing data sets.

| Drug | Platform | Sample ID |  |  |  |
| --- | --- | --- | --- | --- | --- |
|  |  | A1 | A2 | A5 | A7 |
| Tetracycline | BGI | - | tetM | tetM | tetM |
|  | Nanopore | - | - | - | tetM |
| Macrolide | BGI | - | ErmB,RlmA(II) | ErmB,RlmA(II) | ErmB |
|  | Nanopore | - | - | ErmB | ErmB |
| Lincosamide | BGI | - | ErmB,RlmA(II) | ErmB,RlmA(II) | ErmB |
|  | Nanopore | - | - | ErmB | ErmB |
| Streptogramin | BGI | - | ErmB | ErmB | ErmB |
|  | Nanopore | - | - | ErmB | ErmB |
| Fluoroquinolone | BGI | - | patB | patB,pmrA | - |
|  | Nanopore | - | - | - | - |

- No results for this kind of drug.

**FIG 3:**
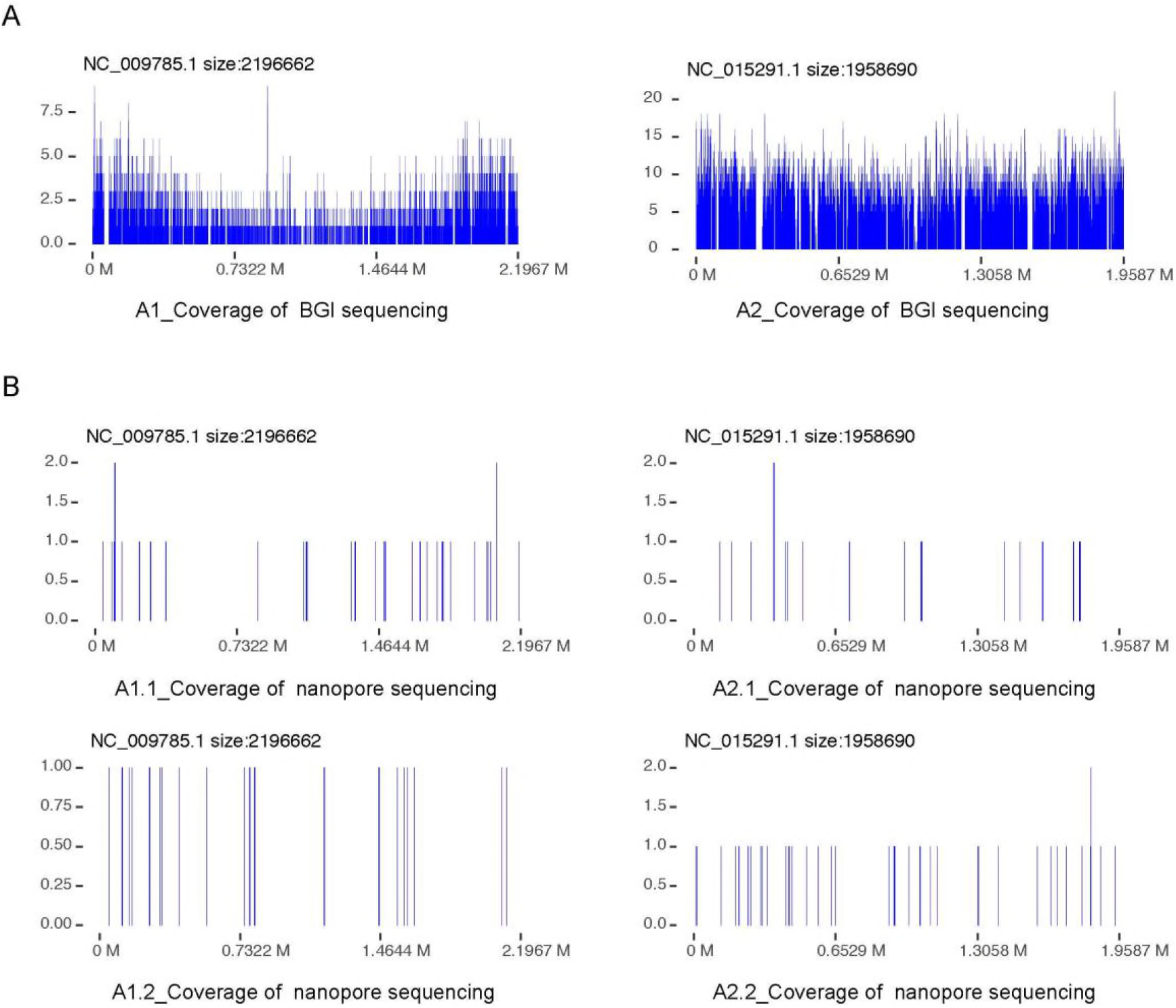
Pathogen coverage of A1 and A2 sequencing data with both NGS and Nanopore MinION platforms. A) the coverage density plot in detected pathogen genome for NGS sequence from BGI platform of A1 and A2 samples; B) the coverage density plot in detected pathogen genome for nanopore sequence from BGI platform of A1 and A2 samples, each sample has two replications. For A1 sample, the detected pathogen is Streptococcus gordonii (NC_009785.1). For A2 sample, the detected pathogen is Streptococcus oralis (NC_015291.1).

### Nanopore sequencing-based metagenomic analysis for IE pathogens detection

To evaluate the application of nanopore sequencing-based metagenomics analysis in IE pathogens detection, DNAs from the seven resected valves were sequenced using the MinION system. In brief, 1.2 μg of genomic DNA from each sample was fragmented with g-TUBE and a library was prepared using the Ligation Sequencing Kit and the Native Barcoding Kit (see details in the Materials and Methods section). The sequencing data generated by the MinION system had a quality score of around 15. This quality score can be influenced by the quality of DNA samples multiplexed in the same flow cell, and high quality multiplexed DNA samples generate larger data with a higher quality score. For every sequencing read, the quality of the first 10 bases can be unstable, with all subsequent bases having a consistent quality score, even for the end bases of an ultra-long read. Reads longer than 1 kb with an average quality score >7, were used in further bioinformatic analyses (see details in the Materials and Methods section).

As a result of metagenomic analysis of the nanopore data, reads of the same IE pathogens were also detected for all samples with NGS (23 and 16 reads of *S. gordonii* for A1.1 and A1.2, 13 and 23 reads of *S. oralis* for A2.1 and A2.2, 68 reads of *C. burnetii* for A3, 2,081 reads of *B. quintana* for A4, 302 reads of *S. oralis* for A5, 42 reads of *S. sanguinis* for A6, and 3,302 reads of *S. anginosus* for A7) (Table 4). Other information such as pathogen coverage, depth, and read length of the nanopore sequencing data were also analyzed (Fig 3B, S1B, and Table S4) with AMR genes of these pathogens detected by the specific bioinformatics pipeline (Fig 2 and Table 3, S3).

**TABLE 4.**
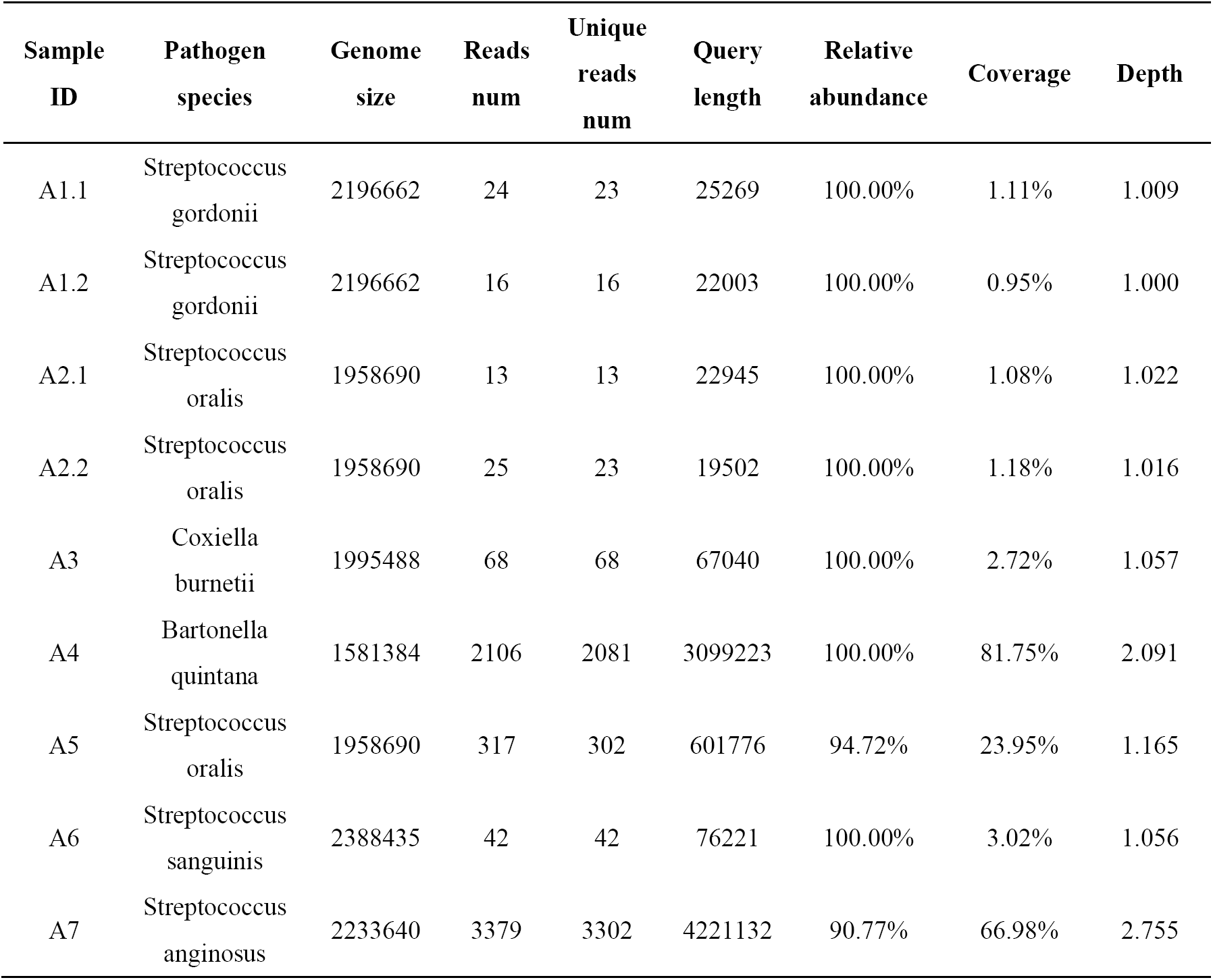
Detail of the results for pathogen species identification from nanopore data with seven samples.

| Sample ID | Pathogen species | Genome size | Reads num | Unique reads num | Query length | Relative abundance | Coverage | Depth |
| --- | --- | --- | --- | --- | --- | --- | --- | --- |
| A1.1 | *Streptococcus gordonii* | 2196662 | 24 | 23 | 25269 | 100.00% | 1.11% | 1.009 |
| A1.2 | *Streptococcus gordonii* | 2196662 | 16 | 16 | 22003 | 100.00% | 0.95% | 1.000 |
| A2.1 | *Streptococcus oralis* | 1958690 | 13 | 13 | 22945 | 100.00% | 1.08% | 1.022 |
| A2.2 | *Streptococcus oralis* | 1958690 | 25 | 23 | 19502 | 100.00% | 1.18% | 1.016 |
| A3 | *Coxiella burnetii* | 1995488 | 68 | 68 | 67040 | 100.00% | 2.72% | 1.057 |
| A4 | *Bartonella quintana* | 1581384 | 2106 | 2081 | 3099223 | 100.00% | 81.75% | 2.091 |
| A5 | *Streptococcus oralis* | 1958690 | 317 | 302 | 601776 | 94.72% | 23.95% | 1.165 |
| A6 | *Streptococcus sanguinis* | 2388435 | 42 | 42 | 76221 | 100.00% | 3.02% | 1.056 |
| A7 | *Streptococcus anginosus* | 2233640 | 3379 | 3302 | 4221132 | 90.77% | 66.98% | 2.755 |

As a real-time sequencing platform, data produced by the MinION system can be base-called and analyzed along with sequencing data. Data generation was rapid during the initiation of sequencing, but decreased with time. After 10 h, negative growth of data was noted. The real-time sequencing properties of the MinION device enabled real-time analysis of pathogens detection, and the minimum stable detection time for a pathogen could be altered by using different detection parameters. For example, if the reads detection cutoff was set at two reads, pathogens in all samples could be detected within 1 h (Fig 4 and Table S5).

**FIG 4:**
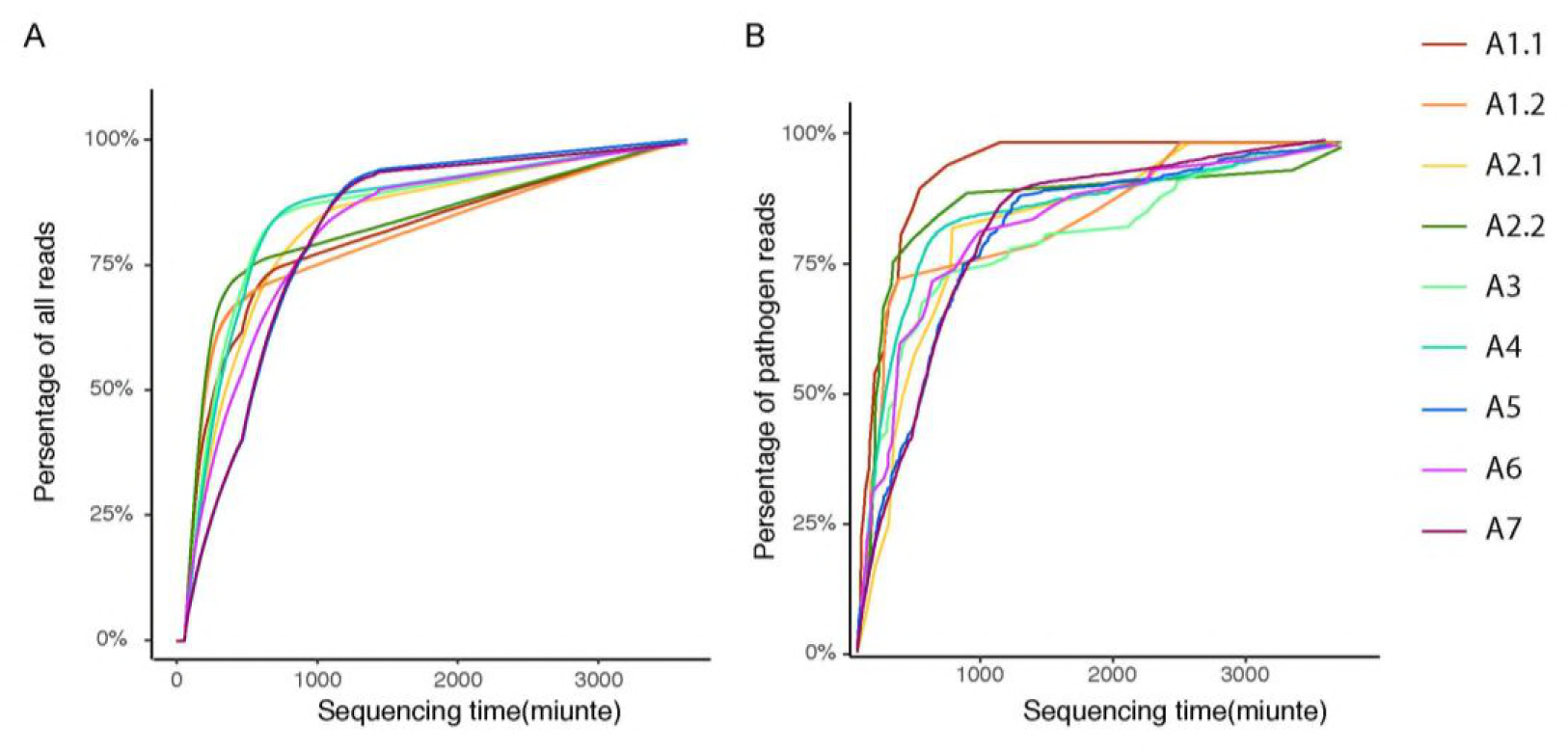
Stable pathogen detection time for different cutoff of reads number in nanopore sequencing data. X axis is the time for sequencing. Y axis is number of reads for detected pathogen in the scale of log2 transfer. Three red dashed lines are the cutoff for pathogen detection, corresponding for difference strict level as two reads, five reads and ten reads. When set two reads as the detection cutoff, all pathogens in samples will be detected within 1 h. Even use a higer cutoff (five reads), all pathogens in samples will be detected within 4 h.

Our results indicated that by integrating real-time nanopore sequencing and appropriate metagenomic bioinformatic approaches, pathogens identification along with the detection of AMR genes could be achieved in cases of culture-negative IE.

## Discussion

Precise diagnosis and effective treatment of IE relies on the rapid and accurate identification of its underlying pathogens. Although blood and valve cultures are the gold standard for IE pathogens detection, blood culture-negative IE can occur in up to 31% of all cases(18).

In this work, we employed both NGS and Oxford Nanopore Technologies MinION nanopore sequencing for pathogens and AMR detection in seven culture-negative IE patients. Our results showed that both methods can reliably identify the causative pathogen in all seven samples in accordance with the results of Sanger sequencing, with the exception of one case in which Sanger sequencing failed (Table 1). Moreover, in the case A2 and A5, Sanger sequencing could only identify bacteria to the genus level whereas NGS and nanopore sequencing-based metagenomics analysis could further classify bacteria to the species level.

Both the NGS and nanopore sequencing results were in agreement in terms of the top enriched species across all samples; however, the remaining species identified were not concordant between the two methods. The NGS results identified a significantly higher number of different bacteria in each sample (Tables S6 and S7). The difference in the amount of sequencing data generated from these two sequencing platforms might contribute to this observation, with a total of 46 Gb of data generated by BGI and only 15 Gb of data generated by MinION for all seven IE samples. Many species identified using the NGS short-reads were of the same genus (Table S6). For example, all nine species detected in A1 belonged to the genus *Streptococcus*, and all 11 species in A2 also belonged to *Streptococcus*. Therefore, we concluded that the long-reads generated by nanopore sequencing increased the specificity of species identification, whereas short-reads generated by NGS had lower resolution within highly homologous species.

For AMR analysis, the extensiveness of pathogen genome coverage was critical. AMR-related genes accounted for only about 1% of the bacterial genome, so broader coverage meant a higher chance of detection. The BGI NGS platform had a much higher data output than the MinION system, resulting in more comprehensive pathogen genome coverage. Therefore, more AMR features were detected using NGS sequencing compared with nanopore sequencing in our study. In terms of the AMR genes detected by both platforms, the NGS results were supported by a significantly higher depth of coverage, which improved the confidence associated with the conclusions drawn from these data. However, the short-reads generated by NGS limited the ability to deduce the origin of AMR genes, i.e. it was not possible to determine the identity of the bacteria carrying a particular AMR feature. If a comparable amount of data can be generated on the nanopore sequencing platform, it offers the advantage of long-reads, which would aid the detection of AMR gene origins. One challenge of AMR detection is to tag the AMR genes to specific microbe because of the high homology of one AMR gene from different species. Sequencing method with longer reads and bigger data volume will favor this goal. In most culture-negative cases, clinicians may have to rely on trial and error during treatment, whereas metagenomic methods can provide pathogens and AMR information, helping to guide clinical drug usage. However, it may be necessary to construct clinic-specific AMR libraries to aid the detection of AMR features.

A few other challenges were observed when analyzing nanopore sequencing data. Sample barcoding is a common practice during library preparation to improve sequencing cost effectiveness by multiplexing samples on one sequencing run. For example, in this study, we multiplexed 3–6 samples for sequencing. Barcode leaking occurred during de-multiplexing when a barcode was misidentified due to a sequencing error. Although barcode leaking is a common problem shared by both NGS and nanopore sequencing platforms, it was much more apparent in the nanopore sequencing results due to its lower sequencing accuracy (advertised base call accuracy of 99.9% for NGS versus 93% for nanopore 1D sequencing). Therefore, to eliminate the possibility of sample cross-contamination on the nanopore sequencing platform, sample multiplexing is not recommended, especially when analyzing clinical samples. The ideal solution in clinical settings is to sequence only one sample per flow cell; this not only avoids contamination but also addresses the clinical point-of-care turnaround time by circumventing the need to batch samples.

Another major challenge in the metagenomic analysis of clinical samples is the high percentage of host genome. More than 95% of sequencing data mapped to the host (human) genome in most IE samples (Table S8), which translates to a huge waste of sequencing data; only approximately 5% of the total sequencing data is actually useable in pathogens identification and AMR detection. Development of appropriate host depletion methods before library preparation will be critical to resolve this problem and increase the percentage of useful sequencing data while maintaining the same amount of total sequencing output, thereby improving detection sensitivity.

In conclusion, the advantages of NGS included low cost, large data volume, and high accuracy rate. In metagenomic analysis, a higher sequencing output correlated with increased sensitivity in pathogens identification and increased confidence in AMR detection. However, the short read-length of NGS was a limiting factor for species identification. For Oxford Nanopore Technologies MinION sequencing, higher cost and lower sequencing data output were limitations in clinical application. However, its unique physical properties and technical features were promising in terms of clinical point-of-care applications. The small size of the device, simple library preparation workflow, real-time sequencing data generation and analysis, and most importantly, long read-length, provided higher accuracy in terms of species identification and AMR linkage.

Our results indicated that the MinION device-based unbiased metagenomic detection of IE pathogens from clinical samples could be performed with a sample-to-answer turnaround time of <1 h if two reads were used as the cutoff and <4 h if five reads were used as the cutoff for species identification. Furthermore, real-time bioinformatic analysis was feasible using nanopore sequencing. All of these features indicated the promising clinical applications of nanopore sequencing-based metagenomic analysis, which were not limited to IE pathogens detection.

Compared with conventional clinical methods, there were some advantages of NGS and nanopore sequencing metagenomic analysis in detecting microorganisms of IE. First, metagenomics analysis could detect unculturable pathogens and overcome the limitations of conventional culture-based methods. Second, metagenomics analysis could detect different types of microorganisms including bacteria, viruses and fungi, whereas 16S rRNA sequencing was limited to screen for bacteria.

Although there are some reports that used NGS-based metagenomic analysis to identify the causative pathogens in culture-negative IE cases^9^, few of these evaluated the usefulness of this new method in AMR gene detection. In this research, we demonstrated that both NGS and nanopore sequencing-based metagenomic analysis could be applied to identify the causative pathogens of IE, thereby providing a valuable, supplemental tool for clinical diagnosis, especially in culture-negative cases. However, before applying metagenomics analysis to clinical microorganism detection, further studies are required to optimize protocols for sample processing, sequencing and bioinformatics analysis.

## Acknowledgements

This work was financially supported by Innovation Project for Medicine and Health Science and Technology from the Chinese Academy of Medical Sciences (Research Project Number:2016-I2M-1-016).

## Author Contributions

Z.Z, Shela Lee and F.W. conceived the idea; J.C., H.H. and W.C. designed the experiments; J.C., H.H., K.W., S.Z., C.C. and Q.C. performed experiments and Y.K. and W.F. analyzed data; J.C. and Q.Z. collected clinical samples. H.H., J.C. and Y.K. wrote the manuscript; Z.Z, Shela Lee, F.W. and A.F. revised the manuscript. All authors read and approved the final manuscript.

